# Mutants libraries reveal negative design shielding proteins from mis-assembly and re-localization in cells

**DOI:** 10.1101/2021.01.20.427404

**Authors:** Hector Garcia Seisdedos, Tal Levin, Gal Shapira, Saskia Freud, Emmanuel Levy

## Abstract

Understanding the molecular consequences of mutations in proteins is essential to map genotypes to phenotypes and interpret the increasing wealth of genomic data. While mutations are known to disrupt protein structure and function, their potential to create new structures and localization phenotypes has not yet been mapped to a sequence space. To map this relationship, we employed two homo-oligomeric protein complexes where the internal symmetry exacerbates the impact of mutations. We mutagenized three surface residues of each complex and monitored the mutations’ effect on localization and assembly phenotypes in yeast cells. While surface mutations are classically viewed as benign, our analysis of several hundred mutants revealed they often trigger three main phenotypes in these proteins: nuclear localization, the formation of puncta, and fibers. Strikingly, more than 50% of random mutants induced one of these phenotypes in both complexes. Analyzing the mutant’s sequences showed that surface stickiness and net charge are two key physicochemical properties associated with these changes. In one complex, more than 60% of mutants self-assembled into fibers. Such a high frequency is explained by negative design: charged residues shield the complex from misassembly, and the sole removal of the charges induces its assembly. A subsequent analysis of several other complexes targeted with alanine mutations suggested that negative design against mis-assembly and mislocalization is common. These results highlight that minimal perturbations in protein surfaces’ physicochemical properties can frequently drive assembly and localization changes in a cellular context.

## INTRODUCTION

Understanding genotype to phenotype relationships is crucial to predict the molecular consequences of mutations (1). At the protein level, alanine scans have revealed how individual residues contribute to protein function, stability, and binding affinity (2–4). More recently, systematic mappings have been widely used to connect sequence variability to changes in protein structure (5, 6), stability (7–9), solubility (10), and functionality (2, 11–14). Similar efforts have been made to map the impact of mutations in protein-ligand (15, 16) and protein-protein interactions (17–21).

However, mutations can impact proteins beyond their stability, function, or existing interactions with specific partners or ligands. Sequences can also encode how proteins distribute spatially in cells, either by addressing them to membrane-bound compartments (22) or by inducing their self-assembly into large repetitive structures (23–27) and membrane-less compartments (28, 29). While changes in protein self-assembly and localization can serve a functional purpose in adaptation (30–36), they can also lead to disease (37). For example, the mis-assembly of hemoglobin and γD-crystallin cause sickle-cell disease and cataracts, respectively (38, 39). The mislocalization of nuclear proteins TDP-43 and FUS in the cytosol is associated with ALS disease (40, 41), and the mislocalization of Ataxin-3 to the nucleus has been implicated in SCA3 disease (42). It is therefore critical to characterize principles by which mutations can trigger such mis-assembly and mislocalization.

Symmetry is frequent in proteins (37, 43) and is a crucial property promoting their self-assembly into high-order structures (44–50). Indeed, a strong enrichment in symmetric homo-oligomers among natural filament-forming proteins has been reported (37). Previous work has also shown that point mutations to two hydrophobic amino acids - leucine and tyrosine - frequently led symmetric homo-oligomers to assemble into high-order assemblies. However, whether other types of amino acids would display a similar potential, whether they would do so often, and whether additional phenotypes of assembly and localization could emerge upon mutation remains unknown.

Here, we assess mutations’ potential to trigger such changes in protein assembly and localization *in vivo*. We targeted two homo-oligomeric protein complexes and randomly mutated three neighboring residues at the surface of each complex. We expressed the mutants fused to a fluorescent protein to track their spatial distribution in yeast cells. We found that a vast sequence space led to changes in protein assembly and localization in both proteins, with three predominant phenotypes: nuclear localization, the formation of filaments, and the formation of puncta. Sequencing of the mutants revealed that increasing surface stickiness frequently promoted nuclear localization in one of the two proteins. Surprisingly, in the other protein, a loss of negatively charged residues was sufficient to trigger protein self-assembly, with fibers frequently forming regardless of the type of mutation, including alanine and glycine. We also observe that five out of ten complexes undergo mis-assembly or a change in cellular localization when surface charges are mutated to alanine, implying that negative design against mis-assembly and mislocalization is common among symmetric homo-oligomers.

## RESULTS

### A plasmid library of surface mutants shows frequent self-assembly and nuclear localization phenotypes

We sought to characterize how frequently mutations can trigger new protein self-assemblies, change protein localization, and identify which types of mutations are most likely to do so. We initially focused on a homo-octameric dipeptidase from *E. coli,* hereafter referred to by its PDB code “1pok” (Figure 1A). To track the self-assembly and localization of the protein in cells, we fused the subunit forming the octamer to a yellow fluorescent protein (YFP, further details of the constructs are provided in Supplementary Table 1). We then created a plasmid library by site-saturation mutagenesis of three solvent-exposed residues (E239/E243/K247) located in an alpha-helix (Figure 1A). We specifically targeted those residues because a triple leucine mutant at these positions was previously observed to form fibers. We transformed the plasmid library in yeast cells and imaged them by fluorescence microscopy (Figure 1B). While the wild-type protein exhibited a homogeneous and cytosolic localization, the mutants frequently formed micrometer-long filaments, puncta, or localized to the nucleus (Figure 1A & B).

**Figure 1.**
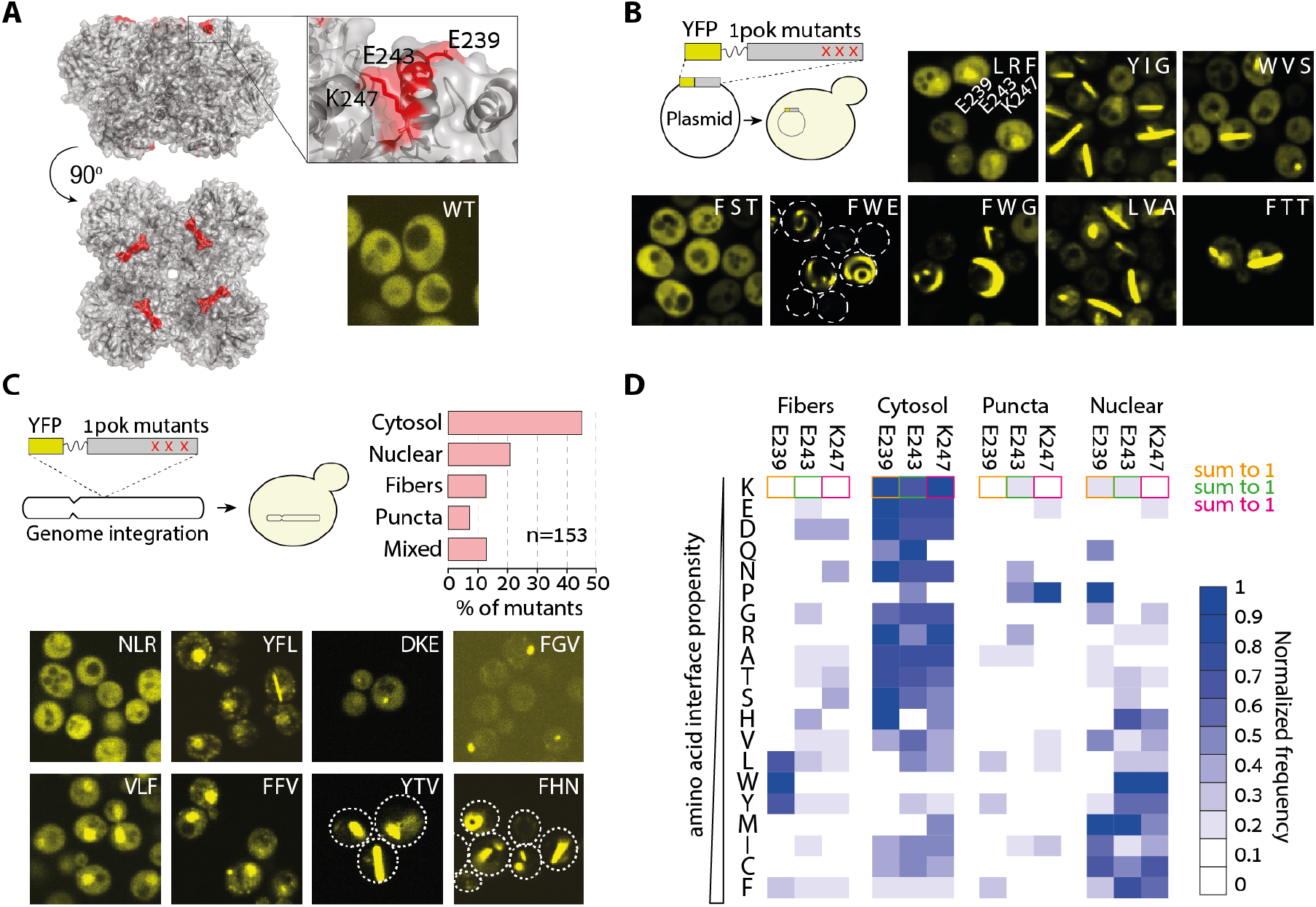
A mutant library reveals a wide diversity of localization phenotypes. **A**. Structure of an octameric protein complex from E. *coli* (PDB code 1POK). Expression of the wild-type homomer fused to a YFP exhibited cytosolic localization. **B**. We created a plasmid library with random mutations in three solvent-exposed residues located at the apex of 1pok and transformed the library into yeast cells. Fluorescence microscopy images of cells expressing representative phenotypes are shown. The identity of the mutations for the residues 239/243/247 is overlaid onto each image. **C.** A different mutant library was created using a genome integration cassette. Subsequent transformation of the library into yeast cells gave genomically integrated mutants. Cells expressing the mutants showed four main localization phenotypes with frequencies given in the barplot. Below we show fluorescence microscopy images of yeast cells displaying eight representative phenotypes. The mutations at residues 239/243/247 are overlaid onto each image. **D.** Frequency of amino-acid at each position (indicated on top) normalized across phenotypes. Amino acids observed in mutants (y-axis, left) are ordered by increasing interaction propensity or “stickiness” (51). The sum of frequencies for a given mutation equals 1.

We subsequently PCR-amplified the regions harboring the mutations to sequence them and relate the various phenotypes to specific amino acid identities. However, sequencing showed that a majority of clones were co-transformants of up to four different plasmids. Consequently, isolation of cells harboring a unique plasmid required multiple streaking steps, making it impossible to evaluate the different phenotypes’ frequencies and relate them to genotypes. Nevertheless, sequencing twenty isolated mutants suggested that filaments and nuclear localization were a frequent outcome of increasing the protein’s surface hydrophobicity (Figure 1B). Notably, one mutant (F/W/E) showed a phenotype with curved filaments growing along the plasma membrane and the nuclear envelope.

### Genome-integrated mutant libraries allow quantifying phenotype frequencies

To overcome the problem of co-transformation with multiple plasmids, we created a new library based on a cassette suitable for genome integration. With this library, each transformant carried a unique set of mutations. In agreement with our initial results, fluorescence microscopy of 220 isolated mutants revealed the frequent formation of fibers and puncta as well as nuclear localization (Supplementary Table 2, Montage 1). Sequencing enabled us to relate the phenotype of each mutant to the underlying mutations. After excluding strains containing deletions, insertions, and stop-codons, we obtained 153 unique variants exhibiting four main phenotypes (Figure 1C): 20.9% displayed nuclear localization, 13.1%, and 7.2% showed fibers and puncta, respectively, and 45.7% of the mutants remained cytosolic like the wild-type sequence. In addition, 13.1% of mutants showed a combination of phenotypes, such as cytosolic localization with a rare presence of puncta or a combination of both fibers and puncta.

We next examined whether specific mutations were associated with particular phenotypes. Given a mutated position (e.g., E239), we calculated the frequency of a target amino acid (e.g., mutation to L) for each of the four main phenotypes, excluding mutants showing mixed phenotypes. For example, out of 14 mutants harboring a leucine at position E239, nine formed fibers, four formed puncta, one was cytosolic, and none exhibited nuclear localization.

Thus, the normalized frequency of leucine-associated-fibers at position E239 is 9/14=0.64. These normalized frequencies for the four phenotypes reveal that fiber-formation is predominantly associated with a mutation to L, W, or Y at position 239 (Figure 1D). The cytosolic phenotype was enriched in mutations to polar amino acids and was favored by mutations to amino acids with low interaction propensity (i.e., non-sticky amino acids as defined previously (51)). Interestingly, nuclear localization appears driven by interaction propensity, in agreement with a recent report (52). Lastly, puncta formation did not show a strong association with a specific set of amino acids except for a large proportion of mutations to proline, suggesting that local unfolding of the alpha-helix might partially explain this phenotype (Figure 1D). Thus, considering the protein 1pok, mutations readily induced changes in its localization and self-assembly phenotypes, and the physicochemical properties of the mutated residues were associated with these changes. Notably, sticky amino acids promoted both new self-assemblies and nuclear localization.

To assess whether mutations could reproducibly induce new assembly and localization phenotypes, we targeted a different protein (Figure 2A): A homo-decameric ketopantoate methyltransferase from *E. coli*, hereinafter referred to by its PDB identifier 1m3u (Figure 2B). Likewise, we created a library of mutants at three solvent-exposed residues located in an alpha-helix. We picked 400 strains, of which 237 carried a unique sequence without stop codons, insertions, or deletions. Phenotypes of the mutants fell into the same four main groups, with a staggering 63.6% of mutants forming fibers, 2.5% localizing to the nucleus, 6.4% forming puncta, 9.7% showing mixed localizations, and only 17.8% remaining cytosolic (Figure 2B & C). The heatmap of mutation’s normalized-frequency for these four phenotypes revealed a different picture than that of 1pok: all types of amino acids promoted fiber formation (Figure 2D). Similarly, the cytosol, puncta, and nuclear localization groups did not exhibit a clear association with a specific class of amino acids (Figure 2D).

**Figure 2.**
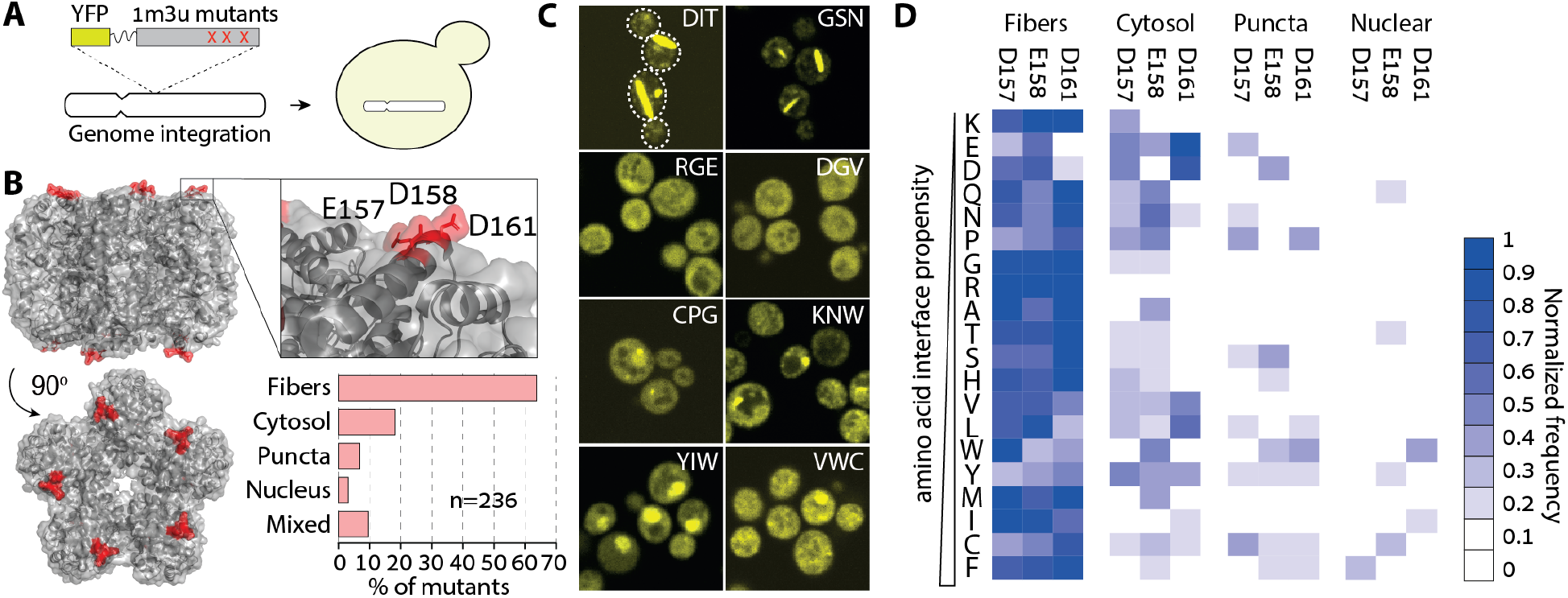
*E. coli* Ketopantoate transferase shows a striking tendency for fiber-formation upon mutation. **A.** We created a YFP-fused 1m3u mutant library in a cassette for genome integration and transformed it into yeast cells. **B.** Structure of the decameric ketopantoate methyltransferase from *E. coli* (PDB code 1M3U). The mutant library targeted three solvent-exposed residues in an alpha helix located at the apex of the complex. Upon expression in yeast cells, mutants displayed four main phenotypes: cytosolic, nuclear, puncta and fiber, with the latter found in >60% of the mutants. **C.** Fluorescence microscopy images of yeast cells expressing eight representative mutants. The identity of mutated residues at positions 157/158/161 is indicated within each image. **D.** Heatmap showing the normalized frequency of mutations of each amino acid type across the four phenotypes.

Overall, both mutant libraries imply that self-assembly and nuclear localization can frequently emerge by surface mutations in homo-oligomers. Indeed, less than 50% of random mutants remained cytosolic in both libraries.

### Stickiness and charge are the main determinants modulating protein assembly and nuclear localization at the level of single cells

We have seen so far that mutations in both homo-oligomers frequently triggered their self-assembly and nuclear localization. However, the relationship between physicochemical changes introduced by the mutations and changes in assembly and localization appeared different for the two proteins studied, which motivates a more quantitative analysis of this genotype-phenotype relationship. Notably, a significant number of strains showed incomplete penetrance of a particular phenotype or even mixed phenotypes. For example, while some strains displayed fibers in most of the cells, in other strains only a small percentage of cells contained fibers (Figure S1).

To capture these phenotypic differences in a quantitative manner, we automatically analyzed the fluorescence properties within single cells and assigned each cell to one or more of the four possible phenotypic categories (Figure S2). This approach enabled us to calculate the fraction cells exhibiting a particular phenotype, i.e., a phenotype’s penetrance, for each mutant strain. The penetrance distribution for each phenotype is shown in Figure 3A. We observe, for example, that fibers are identified in 1.9%, 18.6%, and 79.6% of cells for the 1pok mutants L/G/I, L/V/L, and Y/G/L, respectively (with each triplet corresponding to the positions E239/E243/K247).

**Figure 3.**
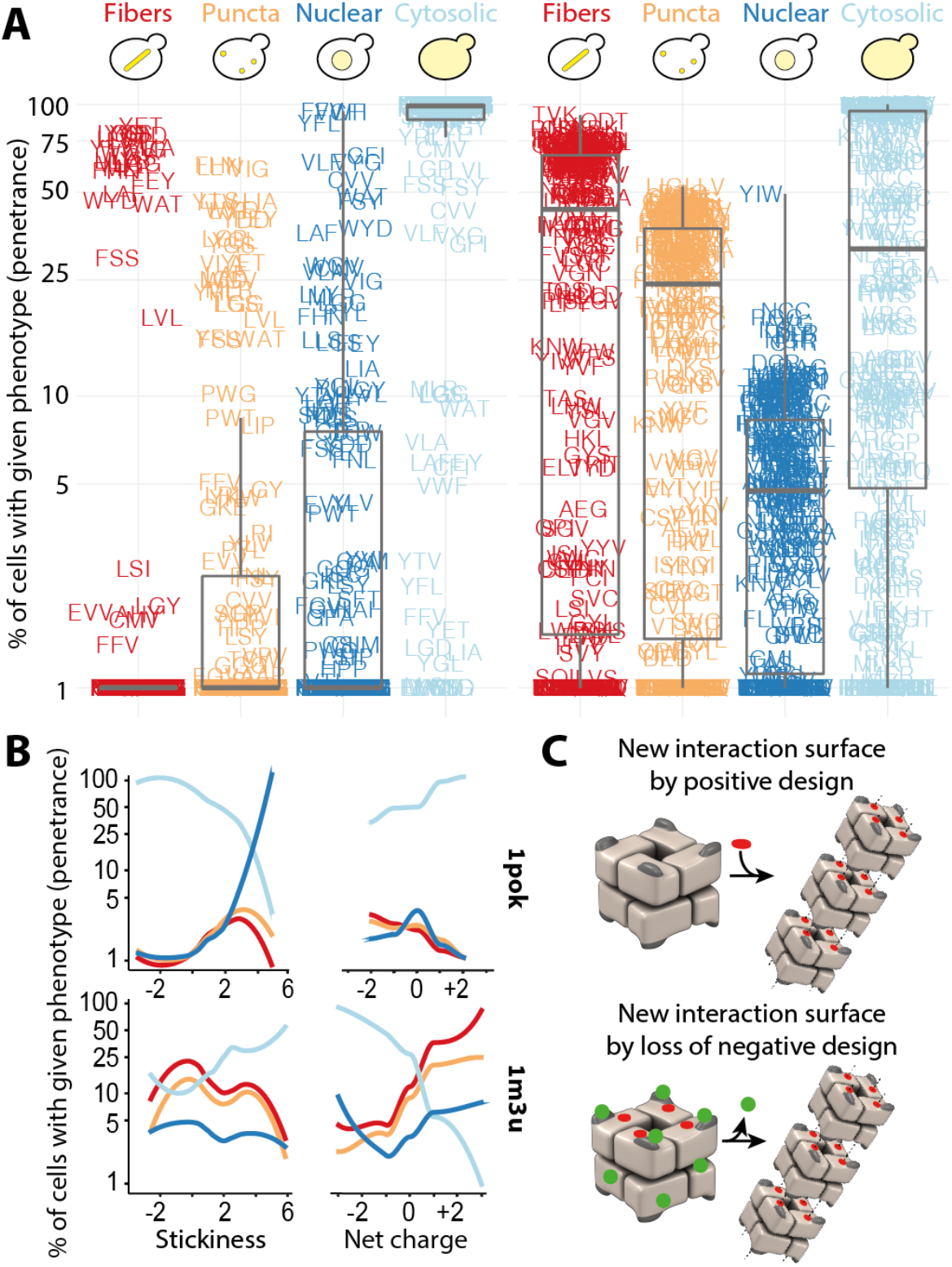
Physico-chemical properties of AA trigger different responses in 1pok and 1m3u. **A** Percentage of cells of a given mutant of 1pok and 1m3u displaying fibers, puncta, cytosolic and nuclear localization (phenotype penetrance). **B** Phenotype penetrance as a function of the summed interaction propensity and net charge of the mutated amino acids for 1pok and 1m3u. **C** Phenotypic changes require an increased stickiness in the case of 1pok, whereas the sole removal of charge appears sufficient to induce phenotypic changes in 1m3u, suggesting that charges represent a feature of “negative design” in this complex.

We analyzed the association between the penetrance of each phenotype and two physicochemical features of the mutated amino acids: their summed interaction propensity and their net charge (Figures 3B & S3). Confirming our initial observation, both proteins exhibited strikingly different patterns. On the one hand, the summed interaction propensity of the triplet was a strong predictor of cytosolic localization in the case of 1pok: 27 and 22 mutants showed an interaction propensity score <0 or >3, and among them, 99% and 46% of cells showed a cytosolic localization, respectively (p=7e-6, Welch’s t-test). In contrast, considering 1m3u, 38, and 37 mutants have an interaction propensity score below 0 or above 3, and these exhibit a similar fraction of cells with cytosolic localization (36.6% and 49.8% respectively, p=0.17). On the other hand, the cytosolic localization of 1m3u was well explained by the net charge of the mutations: among 28 and 62 mutants with a net charge <−1 or >+1, the fraction of cells showing a cytosolic localization was significantly different (73% and 21% respectively, p=9.7e-8). In contrast, the net charge failed to predict cytosolic localization for 1pok. Using the same charge criteria, we find 24 and 26 mutants among which cytosolic localization occurs in 79% and 94% of cells, respectively (p=0.1).

We extended this correlation analysis by considering 20 additional amino-acid features, including secondary structure formation propensity, size, hydrophobicity, etc. (Figure S4, Supplementary Table 4). This analysis confirmed that 1pok and 1m3u respond differently to specific physicochemical changes. While interaction propensity, hydrophobicity, and aromaticity of the residues were the main properties triggering changes in 1pok localization and assembly, 1m3u was primarily impacted by the net charge, as we saw before. We carried out a partial correlation analysis to identify these properties’ independent contributions (Figure S5). This analysis also confirmed the picture sketched above, whereby stickiness is the primary determinant associated with the penetrance of all four phenotypes in 1pok. In contrast, the net charge of the mutations was the main determinant in the case of 1m3u.

Altogether, these results reveal that interaction propensity and charges are central modulators for self-assembly and nuclear localization in these two proteins. These results also highlight that changes in such properties on protein surfaces can trigger different responses in symmetric proteins. For some proteins as 1pok, self-assembly phenotypes may arise as a response to increments in surface hydrophobicity or stickiness, a change that can be regarded as positive design in the context of protein interactions. For other proteins like 1m3u, neutralizing surface charges triggers new phenotypes. Remarkably, in this context, the target amino acid’s identity does not appear particularly important, indicating that the sole removal of charges drives new phenotypes, particularly self-assembly into fibers in this case. This observation points to charged residues at the surface of 1m3u acting as “negative design” elements (Figure 3C), which we examine next.

### Charged gatekeeper residues protect against uncontrolled assembly and re-localization in vivo

The desolvation of hydrophobic residues that accompanies their burial at protein interfaces is energetically favorable (53). For this reason, it is expected that mutations to hydrophobic amino acids are more likely to trigger new interaction sites than mutations to hydrophilic ones. For instance, in sickle cell disease, a charged and polar glutamic acid is mutated to a hydrophobic valine in hemoglobin’s beta chain, inducing its assembly into filaments (38). This idea is consistent with the mutant library of 1pok, whereby self-assembly is associated with positive design in the form of a hydrophobic amino acid at position 239. In stark contrast, the self-assembly of 1m3u into fibers and puncta occurred not only in response to mutations towards hydrophobic residues but also with polar and charged residues. For example, mutants N/S/S, T/D/R, and R/Q/T (at positions D157/D158/D161) consisted exclusively of polar residues and were among the fiber-forming variants (Supplementary Table 2). These mutants suggest that the sole elimination of the negatively charged residues suffices to create a new self-interaction and induce the self-assembly of 1m3u into fibers. This idea is reminiscent of negative design (54–59), where gatekeeper residues prevent proteins from folding into non-native conformations (60, 61) and from engaging in non-native interactions (54). Indeed, removing elements of negative design can promote intermolecular contacts and protein crystallization (62).

To test whether disabling the negatively charged residues sufficed to trigger new self-interactions, we created specific triple mutants to alanine. Interestingly, the mutations drove the formation of fibers and puncta in 1m3u, whereas 1pok remained soluble (Figure 4A). We observed a similar outcome when examining the effect of triple glycine mutants of 1pok and 1m3u that existed in the mutant libraries (Supplementary Table 2). Taken together, these results suggest that self-assembly of 1pok into fibers requires positive design, whereas removal of charges at the surface of 1m3u is sufficient to trigger its assembly into fibers.

**Figure 4.**
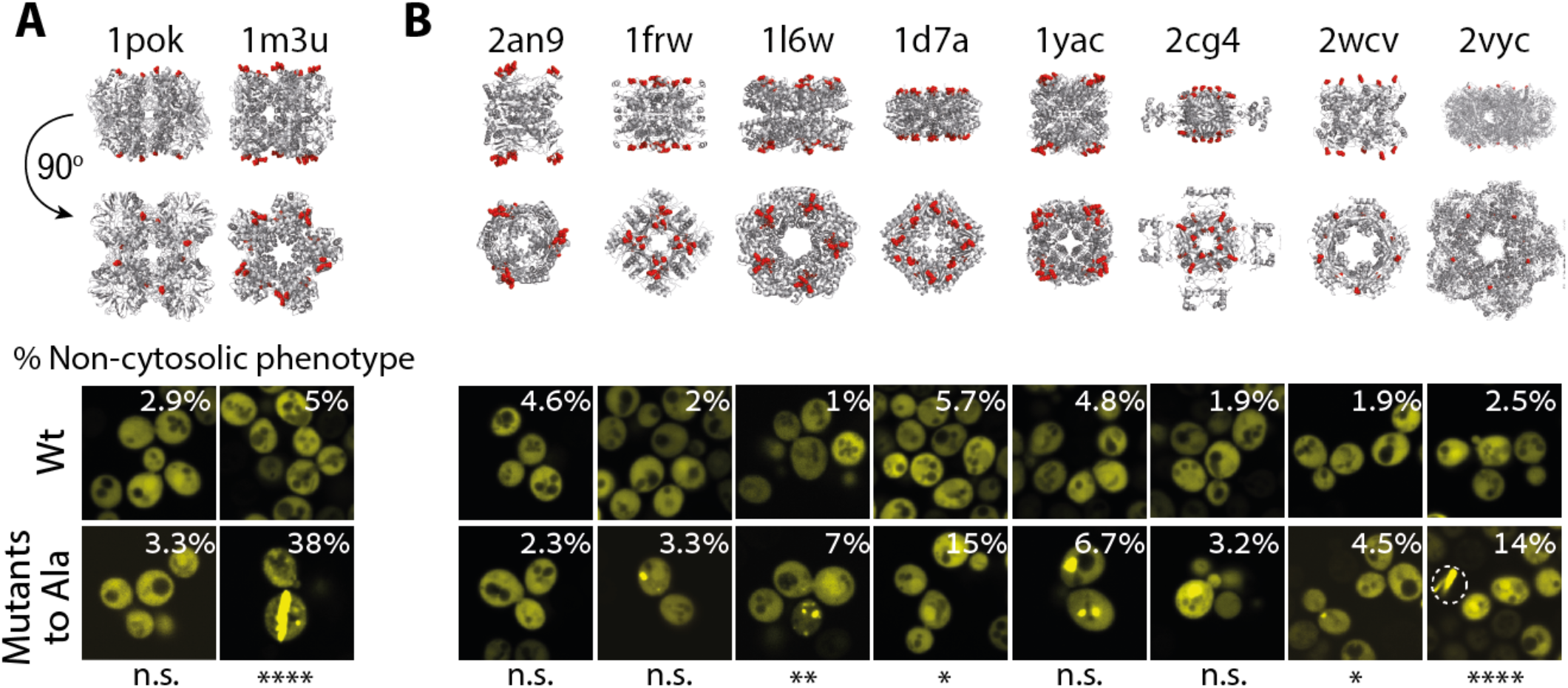
Negative design against self-assembly is common in symmetric proteins. **A.** Fluorescence microscopy images of yeast cells expressing the wild-type and the alanine mutants in 1pok and 1m3u. **B.** Fluorescence microscopy images of yeast cells expressing eight additional symmetric complexes and their alanine mutant variants. The percentage of cells with a non-cytosolic phenotype for each strain is overlaid onto the image.

To investigate this idea further, we created alanine mutants for eight additional homomers, which we refer to by their PDB code (Figure 4B, Supplementary Table 3). We introduced two to four mutations at positions previously shown to trigger the formation of fibers when mutated to leucine or tyrosine (44). The mutations to alanine increased the penetrance of non-cytosolic phenotypes in seven out of the eight complexes. Triggering more frequent puncta in 1l6w, 1frw, 1yac, 2wcv, fibers in 2vyc, and nuclear localization in 1d7a and 2cg4 (Figure 4B). Thus, the sole neutralization of surface charges can, by itself, often trigger new protein assemblies and change a protein’s subcellular localization.

## DISCUSSION AND CONCLUSIONS

Here, we demonstrate that mutations at the surface of two symmetric complexes frequently trigger their self-assembly into puncta and fibers, as well as changes in their subcellular localization. Furthermore, we report two distinct mutational pathways driving these changes: One involves mutations increasing surface stickiness, another the sole removal of surface charges. Thus, while the first route relies on positive design, the second involves neutralizing negative design elements. This result prompted us to analyze eight additional symmetric complexes and ask whether mutating surface charges to alanine could suffice to trigger new assembly and localization phenotypes. We detected such changes in nine out of ten protein complexes investigated, and for five, the changes were statistically significant. This observation confirms experimentally that negative design is frequent among symmetric protein complexes.

In this work, we focused on the effect of mutations at patches located at the apex of the quaternary structure (Figure 4). In previous work, we predicted that these regions are subject to negative design (44, 49), and here we experimentally validated this prediction. Future work will be needed to identify whether mutations in other parts of the structure (i.e., farther from the “apex”) also frequently trigger changes in protein assembly and subcellular localization phenotypes.

The two mutational pathways we observed also occur among natural protein assemblies. Indeed, numerous protein-forming filaments are stabilized by the interaction of specific amino acids that are typically hydrophobic or aromatic. For example, glutamic acid to valine mutation is causing hemoglobin to form filaments and leads to sickle cell disease (38). In another example, the polymerization of yeast hexokinase into filaments occurs when a tyrosine inserts in a hydrophobic pocket due to a ligand-triggered conformational change (30). Similar interactions are responsible for the filament formation of human IMPDH (63), the *E. coli* and human CTPS (64), and for the drug-induced polymerization of the oncogenic transcription factor BCL6 (65).

Interestingly, the increase in self-assembly propensity seen in quiescent cells is associated with changes in pH (34, 66) and might be reminiscent of the negative design principle observed in this work. Indeed, a decrease in pH may neutralize surface charges and unlock cryptic interaction sites, promoting self-assembly in those conditions. Interestingly, the oxidative damage associated with aging cells promotes aggregation through charge neutralization (67), and a similar mechanism could trigger protein assembly. Future experiments aimed to map the sequence-assembly relation in different growth conditions or at different pH will help to single out sequence determinants for such condition-dependent protein assembly.

A striking number of mutants of the 1pok and 1m3u libraries and some of the alanine mutants of two additional complexes (1d7a and 2cg4) exhibited nuclear localization. The nuclear pore complex (NPC) has a diffusion barrier composed of intrinsically disordered proteins rich in phenylalanine and glycine (FG domains). It acts as a molecular sieve with a passive-diffusion size-limit of about 40 kDa. Larger macromolecules do not efficiently cross the diffusion barrier and must be ferried through by nuclear transporters (68). All the protein complexes used in this work considerably exceed the limit of passive diffusion (Supplementary Tables 1 and 3). Specifically, 1m3u and 1pok have molecular weights in the range of 280 kDa and 320 kDa, respectively. Since they lack an NLS signal, they should not be recognized by nuclear transporters, and therefore, they should be excluded from the nucleus. However, in agreement with previous reports (52, 69), we found that surface hydrophobicity promotes the nuclear localization of 1pok despite its large size. At the same time, the nuclear localization of 1m3u is not driven by an increase of surface stickiness but rather by positive charges. Interestingly, the fact that mutations promoting nuclear localization are of a different type for both complexes and promote their respective self-assembly in fibers and puncta suggests that the nuclear localization we observe is linked to the formation of high-order assemblies. The dysregulation of nucleo-cytoplasmic localization is implicated in neurodegenerative disorders as well as cancer (70). Our results imply that such dysregulation of localization can occur through non-canonical pathways that will need to be characterized in future work.

This work paves the way to understand how changes in a protein’s surface chemistry can impact its spatial distribution in the cell by modulating its self-assembly state and nucleo-cytoplasmic localization. A surprisingly wide sequence space drove assembly and localization changes in the two proteins that we studied, and future work will be needed to generalize these observations to other proteins. In that respect, the fact that proteins are marginally soluble (71) means we could expect such assembly and localization changes to be common. The observations we made also bear implications in terms of proteome function and evolution. While protein localization is regarded as being regulated by linear motifs, changes in protein surface chemistry may drive such changes during evolution, both in health and disease.

## MATERIALS AND METHODS

### Selection of the proteins

The homo-oligomers used in this work were selected based on specific criteria described previously (44). Details of their structure, gene names, PDB accession numbers, number of subunits, symmetry, theoretical isoelectric point, and number of positives and negatives charges are given in Supplementary Tables 1 and 3.

### Cloning procedures, mutagenesis, and gene libraries construction

Genes coding for the ten dihedral homomers used in this work were amplified from the *E. coli* strain K12 and cloned as previously described (44). Additionally, genes encoding the structures 1POK and 1M3U were cloned downstream of the yeast GPD promoter into M3925 plasmids for genome integration (72). Molecular cloning was performed using the PIPE method (73). The ten homomers were fused to a YFP (Venus) (74) spaced by a flexible linker (sequence: GGGGSGGGGS) to track their localization *in vivo*. To introduce alanine mutations, the ten homomers were subjected to site-directed mutagenesis (QuickChange, Agilent). The 1pok episomal gene library was based on the plasmid p413 GPD. The genomically-integrated mutant libraries were based on the plasmid M3925 GPD (trp1::KanMX3). Site saturation mutagenesis of the targeted residues (Supplementary Table 1) was achieved using oligos with randomized bases in the codons targeted for mutagenesis. All residues targeted for mutagenesis were solvent-exposed with >25% accessible surface area. The oligonucleotides used for cloning, mutagenesis, and mutant library construction were ordered from IDT.

### Resolving multiple vector transformants

Sequencing of yeast colonies after transformation of the p413 plasmid-based 1pok gene library revealed that most of them bore up to four different plasmids. To resolve individual sequences, single colonies were diluted in PBS and streaked on agar plates with synthetic complete medium lacking histidine (SC-His). This dilution-streaking procedure was iterated three times for every mutant before they were sequenced.

### Yeast strains and media

The parent yeast strain used in this work was BY4741 (MATa his3Δ1 leu2Δ0 met15Δ0 ura3Δ0). Plasmids p413 encoding for the wild-type proteins, the alanine mutants, and the 1pok gene library were transformed into BY4741 cells using the LiAc method, as described (75). Transformants were grown in SC-His agar plates and were inoculated into 384-well plates containing SC-His with 15% glycerol and stored −80°C after growing to saturation. For the 1pok and 1m3u gene libraries based in the M3925 plasmid, the cassette for genomic integration was amplified as a linear fragment prior to its transformation into yeast cells. Transformants were grown in SC + G418 (300 μg/ml) agar plates, and were also inoculated into 384-well plates, grown in SC + G418 (200 μg/ml) 15% glycerol, and stored at −80°C.

### Microscopy Screenings and sample preparation

A liquid handling robot (Tecan Evo 200) was used to perform plate-to-plate transfers of cells by operating a pintool (FP1 pins, V&P scientific). Cells were inoculated from glycerol stocks into 384-well polypropylene plates containing SC-His (for strains containing p413 plasmids) or SC + G418 (200 μg/ml) (for strains of the genome-integrated gene libraries) and grown until they reached saturation. Then, cells from the saturated cultures were inoculated into Greiner™ 384-well glass-bottom optical imaging plates containing the appropriate selection media for each case. Cells were grown for at least 6 hours until they reached logarithmic growth (an optical density of ~0.5-1). Imaging was performed using an automated Olympus microscope X83 coupled to a spinning disk confocal scanner (Yokogawa W1) with a 60x, 1.35 NA, oil immersion objective (UPLSAPO 60XO, Olympus). Fluorescence excitation was achieved with a 488 nm laser (Toptica, 100mW), emission filter sets were (525/28, Chroma). We recorded 16-bit images for brightfield and GFP channels on a Hamamatsu Flash 4 V2 camera. Hardware autofocus (Olympus Z-Drift Compensation System) was used to maintain the focus during the imaging experiments.

### Image Analysis

Cells were identified, segmented, and their fluorescent signal (median, average, minimum, maximum) as well as additional cell properties were determined using custom algorithms (76, 77) in FIJI (78) and exported as tabulated files.

### Automatic assignment of cell phenotypes

Fibers, puncta, and nuclear localization were identified in each cell independently, in a multistep process: (i) We calculated the maximum and median fluorescence intensity of pixels in a given cell. (ii) Cells where the maximal intensity was below 200 or cells not containing a markedly bright region (maximum/median < 2.5) were assigned as cytosolic. (iii) We identified the largest four regions composed of pixels with an intensity 2.5-fold above the cell median intensity. If such regions of interest (ROI) existed, showed a circularity above 0.4 and an area above 4 pixels, the cell was assigned one or more of the non-cytosolic phenotypes (Fibers, Nuclear or Puncta) based on criteria of the ROI fluorescence and shape, as described in a decision tree (Figure S2B). In those cells, for each ROI found (up to the fourth-largest), a phenotype was assigned to the ROI. Finally, the phenotype of the cell was determined as the union of the phenotypes of all the ROIs. This procedure was optimized based on a dataset of 1834 cells that we annotated manually.

### Data analysis

The 20 numerical scales representing physicochemical and biochemical properties of AAs (Supplementary Table 4) were selected from (http://www.genome.jp/aaindex/), except for interaction propensities scales taken from (79) and (Villegas J *et al.* manuscript in preparation). Data from sequencing was tabulated and aggregated with phenotypic information (Tables S2 and S4). The physicochemical properties of the mutants and their relationship to phenotypic information was analyzed using R.

## Supporting information

Table S2--1pok

## Author Contribution

H.G.S. and E.D.L. designed the experiments. H.G.S., G.S., and S.F. performed the experiments. H.G.S., T.L., E.D.L. analyzed the data, H.G.S. and E.D.L. wrote the manuscript.

## Acknowledgments

We thank Benjamin Dubreuil for helping with the data analysis, and H. Greenblatt for helping with the computer infrastructure. This work was supported by the European Research Council (ERC) under the European Union’s Horizon 2020 research and innovation program (grant agreement No. 819318), by the Israel Science Foundation (grant no. 1452/18), by a research grant from A.-M. Boucher, by research grants from the Estelle Funk Foundation, the Estate of Fannie Sherr, the Estate of Albert Delighter, the Merle S. Cahn Foundation, Mrs. Mildred S. Gosden, the Estate of Elizabeth Wachsman, the Arnold Bortman Family Foundation. H.G.S. received support from the Koshland Foundation.

## REFERENCES

1. P. C. Ng, S. Henikoff, Predicting the effects of amino acid substitutions on protein function. Annu. Rev. Genomics Hum. Genet. 7, 61–80 (2006).

2. D. M. Fowler, et al., High-resolution mapping of protein sequence-function relationships. Nat. Methods 7, 741–746 (2010).

3. B. C. Cunningham, J. A. Wells, High-resolution epitope mapping of hGH-receptor interactions by alanine-scanning mutagenesis. Science 244, 1081–1085 (1989).

4. G. A. Weiss, C. K. Watanabe, A. Zhong, A. Goddard, S. S. Sidhu, Rapid mapping of protein functional epitopes by combinatorial alanine scanning. Proc. Natl. Acad. Sci. U. S. A. 97, 8950–8954 (2000).

5. W. Huang, J. Petrosino, M. Hirsch, P. S. Shenkin, T. Palzkill, Amino acid sequence determinants of beta-lactamase structure and activity. J. Mol. Biol. 258, 688–703 (1996).

6. J. Suckow, et al., Genetic studies of the Lac repressor. XV: 4000 single amino acid substitutions and analysis of the resulting phenotypes on the basis of the protein structure. J. Mol. Biol. 261, 509–523 (1996).

7. G. J. Rocklin, et al., Global analysis of protein folding using massively parallel design, synthesis, and testing. Science 357, 168–175 (2017).

8. K. M. Schlinkmann, et al., Critical features for biosynthesis, stability, and functionality of a G protein-coupled receptor uncovered by all-versus-all mutations. Proc. Natl. Acad. Sci. U. S. A. 109, 9810–9815 (2012).

9. I. Kim, C. R. Miller, D. L. Young, S. Fields, High-throughput analysis of in vivo protein stability. Mol. Cell. Proteomics 12, 3370–3378 (2013).

10. B. Bolognesi, et al., The mutational landscape of a prion-like domain. Nat. Commun. 10, 4162 (2019).

11. H. H. Guo, J. Choe, L. A. Loeb, Protein tolerance to random amino acid change. Proc. Natl. Acad. Sci. U. S. A. 101, 9205–9210 (2004).

12. E. Firnberg, J. W. Labonte, J. J. Gray, M. Ostermeier, A comprehensive, high-resolution map of a gene’s fitness landscape. Mol. Biol. Evol. 31, 1581–1592 (2014).

13. L. Rockah-Shmuel, Á. Tóth-Petróczy, D. S. Tawfik, Systematic Mapping of Protein Mutational Space by Prolonged Drift Reveals the Deleterious Effects of Seemingly Neutral Mutations. PLoS Comput. Biol. 11, e1004421 (2015).

14. P. C. Després, A. K. Dubé, M. Seki, N. Yachie, C. R. Landry, Perturbing proteomes at single residue resolution using base editing. Nat. Commun. 11, 1871 (2020).

15. A. Ernst, et al., Coevolution of PDZ domain-ligand interactions analyzed by high-throughput phage display and deep sequencing. Mol. Biosyst. 6, 1782–1790 (2010).

16. N. D. Taylor, et al., Engineering an allosteric transcription factor to respond to new ligands. Nat. Methods 13, 177–183 (2016).

17. S. S. Sidhu, S. Koide, Phage display for engineering and analyzing protein interaction interfaces. Curr. Opin. Struct. Biol. 17, 481–487 (2007).

18. T. A. Whitehead, et al., Optimization of affinity, specificity and function of designed influenza inhibitors using deep sequencing. Nat. Biotechnol. 30, 543–548 (2012).

19. J. G. Jardine, et al., HIV-1 broadly neutralizing antibody precursor B cells revealed by germline-targeting immunogen. Science 351, 1458–1463 (2016).

20. R. Cohen-Khait, G. Schreiber, Low-stringency selection of TEM1 for BLIP shows interface plasticity and selection for faster binders. Proc. Natl. Acad. Sci. U. S. A. 113, 14982–14987 (2016).

21. G. Diss, B. Lehner, The genetic landscape of a physical interaction. Elife 7 (2018).

22. W.-K. Huh, et al., Global analysis of protein localization in budding yeast. Nature 425, 686–691 (2003).

23. S. Alberti, R. Halfmann, O. King, A. Kapila, S. Lindquist, A systematic survey identifies prions and illuminates sequence features of prionogenic proteins. Cell 137, 146–158 (2009).

24. F. Cid-Samper, et al., An Integrative Study of Protein-RNA Condensates Identifies Scaffolding RNAs and Reveals Players in Fragile X-Associated Tremor/Ataxia Syndrome. Cell Rep. 25, 3422–3434.e7 (2018).

25. R. M. Vernon, et al., Pi-Pi contacts are an overlooked protein feature relevant to phase separation. Elife 7 (2018).

26. J. Wang, et al., A Molecular Grammar Governing the Driving Forces for Phase Separation of Prion-like RNA Binding Proteins. Cell 174, 688–699.e16 (2018).

27. M. P. Hughes, et al., Atomic structures of low-complexity protein segments reveal kinked β sheets that assemble networks. Science 359, 698–701 (2018).

28. A. A. Hyman, C. A. Weber, F. Jülicher, Liquid-liquid phase separation in biology. Annu. Rev. Cell Dev. Biol. 30, 39–58 (2014).

29. E. Gomes, J. Shorter, The molecular language of membraneless organelles. J. Biol. Chem. 294, 7115–7127 (2019).

30. P. R. Stoddard, et al., Polymerization in the actin ATPase clan regulates hexokinase activity in yeast. Science 367, 1039–1042 (2020).

31. R. M. Barry, et al., Large-scale filament formation inhibits the activity of CTP synthetase. Elife 3, e03638 (2014).

32. K. C. Duong-Ly, et al., T cell activation triggers reversible inosine-5′-monophosphate dehydrogenase assembly. Journal of Cell Science 131, jcs223289 (2018).

33. C.-W. Kim, et al., Induced polymerization of mammalian acetyl-CoA carboxylase by MIG12 provides a tertiary level of regulation of fatty acid synthesis. Proc. Natl. Acad. Sci. U. S. A. 107, 9626–9631 (2010).

34. I. Petrovska, et al., Filament formation by metabolic enzymes is a specific adaptation to an advanced state of cellular starvation. Elife (2014) https:/doi.org/10.7554/eLife.02409.

35. R. Narayanaswamy, et al., Widespread reorganization of metabolic enzymes into reversible assemblies upon nutrient starvation. Proc. Natl. Acad. Sci. U. S. A. 106, 10147–10152 (2009).

36. C. Noree, et al., A quantitative screen for metabolic enzyme structures reveals patterns of assembly across the yeast metabolic network. Mol. Biol. Cell 30, 2721–2736 (2019).

37. H. Garcia-Seisdedos, J. A. Villegas, E. D. Levy, Infinite Assembly of Folded Proteins in Evolution, Disease, and Engineering. Angewandte Chemie International Edition 58, 5514–5531 (2019).

38. W. A. Eaton, J. Hofrichter, Sickle cell hemoglobin polymerization. Adv. Protein Chem. 40, 63–279 (1990).

39. J. C. Boatz, M. J. Whitley, M. Li, A. M. Gronenborn, P. C. A. van der Wel, Cataract-associated P23T γD-crystallin retains a native-like fold in amorphous-looking aggregates formed at physiological pH. Nat. Commun. 8, 15137 (2017).

40. S. J. Barmada, et al., Cytoplasmic mislocalization of TDP-43 is toxic to neurons and enhanced by a mutation associated with familial amyotrophic lateral sclerosis. J. Neurosci. 30, 639–649 (2010).

41. T. J. Kwiatkowski Jr, et al., Mutations in the FUS/TLS gene on chromosome 16 cause familial amyotrophic lateral sclerosis. Science 323, 1205–1208 (2009).

42. P. M. A. Antony, et al., Identification and functional dissection of localization signals within ataxin-3. Neurobiol. Dis. 36, 280–292 (2009).

43. J. A. Marsh, S. A. Teichmann, Structure, dynamics, assembly, and evolution of protein complexes. Annu. Rev. Biochem. 84, 551–575 (2015).

44. H. Garcia-Seisdedos, C. Empereur-Mot, N. Elad, E. D. Levy, Proteins evolve on the edge of supramolecular self-assembly. Nature 548, 244–247 (2017).

45. J. E. Padilla, C. Colovos, T. O. Yeates, Nanohedra: using symmetry to design self assembling protein cages, layers, crystals, and filaments. Proc. Natl. Acad. Sci. U. S. A. 98, 2217–2221 (2001).

46. N. P. King, et al., Computational design of self-assembling protein nanomaterials with atomic level accuracy. Science 336, 1171–1174 (2012).

47. Y. Suzuki, et al., Self-assembly of coherently dynamic, auxetic, two-dimensional protein crystals. Nature 533, 369–373 (2016).

48. D. Grueninger, et al., Designed protein-protein association. Science 319, 206–209 (2008).

49. C. Empereur-Mot, H. Garcia-Seisdedos, N. Elad, S. Dey, E. D. Levy, Geometric description of self-interaction potential in symmetric protein complexes. Sci Data 6, 64 (2019).

50. A. J. Ben-Sasson, et al., Design of biologically active binary protein 2D materials. Nature (2021) https:/doi.org/10.1038/s41586-020-03120-8.

51. E. D. Levy, S. De, S. A. Teichmann, Cellular crowding imposes global constraints on the chemistry and evolution of proteomes. Proc. Natl. Acad. Sci. U. S. A. 109, 20461–20466 (2012).

52. S. Frey, et al., Surface Properties Determining Passage Rates of Proteins through Nuclear Pores. Cell 174, 202–217.e9 (2018).

53. C. Chothia, J. Janin, Principles of protein–protein recognition. Nature 256, 705–708 (1975).

54. J. P. K. Doye, A. A. Louis, M. Vendruscolo, Inhibition of protein crystallization by evolutionary negative design. Phys. Biol. 1, P9–13 (2004).

55. S. Pechmann, E. D. Levy, G. G. Tartaglia, M. Vendruscolo, Physicochemical principles that regulate the competition between functional and dysfunctional association of proteins. Proc. Natl. Acad. Sci. U. S. A. 106, 10159–10164 (2009).

56. D. N. Bolon, R. A. Grant, T. A. Baker, R. T. Sauer, Specificity versus stability in computational protein design. Proc. Natl. Acad. Sci. U. S. A. 102, 12724–12729 (2005).

57. I. N. Berezovsky, K. B. Zeldovich, E. I. Shakhnovich, Positive and negative design in stability and thermal adaptation of natural proteins. PLoS Comput. Biol. 3, e52 (2007).

58. O. Noivirt-Brik, A. Horovitz, R. Unger, Trade-off between positive and negative design of protein stability: from lattice models to real proteins. PLoS Comput. Biol. 5, e1000592 (2009).

59. J. S. Richardson, D. C. Richardson, Natural β-sheet proteins use negative design to avoid edge-to-edge aggregation. Proceedings of the (2002).

60. W. F. DeGrado, C. M. Summa, V. Pavone, F. Nastri, A. Lombardi, De novo design and structural characterization of proteins and metalloproteins. Annu. Rev. Biochem. 68, 779–819 (1999).

61. M. H. Hecht, J. S. Richardson, D. C. Richardson, R. C. Ogden, De novo design, expression, and characterization of Felix: a four-helix bundle protein of native-like sequence. Science 249, 884–891 (1990).

62. Z. S. Derewenda, Rational protein crystallization by mutational surface engineering. Structure 12, 529–535 (2004).

63. S. A. Anthony, et al., Reconstituted IMPDH polymers accommodate both catalytically active and inactive conformations. Mol. Biol. Cell (2017) https:/doi.org/10.1091/mbc.E17-04-0263.

64. E. M. Lynch, et al., Human CTP synthase filament structure reveals the active enzyme conformation. Nat. Struct. Mol. Biol. 24, 507–514 (2017).

65. M. Słabicki, et al., Small-molecule-induced polymerization triggers degradation of BCL6. Nature 588, 164–168 (2020).

66. M. C. Munder, et al., A pH-driven transition of the cytoplasm from a fluid- to a solid-like state promotes entry into dormancy. eLife 5 (2016).

67. A. M. R. de Graff, M. J. Hazoglou, K. A. Dill, Highly Charged Proteins: The Achilles’ Heel of Aging Proteomes. Structure 24, 329–336 (2016).

68. D. H. Lin, A. Hoelz, The Structure of the Nuclear Pore Complex (An Update). Annu. Rev. Biochem. 88, 725–783 (2019).

69. B. Naim, D. Zbaida, S. Dagan, R. Kapon, Z. Reich, Cargo surface hydrophobicity is sufficient to overcome the nuclear pore complex selectivity barrier. EMBO J. 28, 2697–2705 (2009).

70. M.-C. Hung, W. Link, Protein localization in disease and therapy. J. Cell Sci. 124, 3381–3392 (2011).

71. G. Vecchi, et al., Proteome-wide observation of the phenomenon of life on the edge of solubility. Proc. Natl. Acad. Sci. U. S. A. 117, 1015–1020 (2020).

72. W. P. Voth, Y. W. Jiang, D. J. Stillman, New “marker swap” plasmids for converting selectable markers on budding yeast gene disruptions and plasmids. Yeast 20, 985–993 (2003).

73. H. E. Klock, E. J. Koesema, M. W. Knuth, S. A. Lesley, Combining the polymerase incomplete primer extension method for cloning and mutagenesis with microscreening to accelerate structural genomics efforts. Proteins 71, 982–994 (2008).

74. T. Nagai, et al., A variant of yellow fluorescent protein with fast and efficient maturation for cell-biological applications. Nat. Biotechnol. 20, 87–90 (2002).

75. M. Knop, et al., Epitope tagging of yeast genes using a PCR-based strategy: more tags and improved practical routines. Yeast 15, 963–972 (1999).

76. O. Matalon, A. Steinberg, E. Sass, J. Hausser, E. D. Levy, Reprogramming protein abundance fluctuations in single cells by degradation. bioRxiv (2018) https:/doi.org/10.1101/260695.

77. M. Heidenreich, et al., Designer protein assemblies with tunable phase diagrams in living cells. Nat. Chem. Biol. 16, 939–945 (2020).

78. J. Schindelin, et al., Fiji: an open-source platform for biological-image analysis. Nat. Methods 9, 676–682 (2012).

79. E. D. Levy, A simple definition of structural regions in proteins and its use in analyzing interface evolution. J. Mol. Biol. 403, 660–670 (2010).

