## Supplementary Figures and Tables S1 and S3 for "Mutants libraries reveal negative design shielding proteins from mis-assembly and re-localization in cells": GarciaSeisdedos_et_al_SUP.pdf

**Supplementary Table 1.** Details of homomers targeted for mutagenesis: PDB code, number of subunits, symmetry, ORF, mutated positions, linker, and fluorescent protein fused.

| PDB code | Gene Name | Ref. | # subunits | MW (kDa, per subunit) | Symmetry | pI wild-type | pI Ala mut | # + charges | # - charges | ORF | Mutated residues | Linker | Fluorophore |
|---|---|---|---|---|---|---|---|---|---|---|---|---|---|
| 1POK | *ladA* | (1) | 8 | 41.1 | D4 | 5.08 | 5.13 | 31 | 46 | Venus-1pok | E239/E243/K247 | GGGGS GGGGS | Venus (2) |
| 1M3U | *panB* | (3) | 10 | 28.2 | D5 | 5.15 | 5.5 | 20 | 30 | 1m3u-Venus | D157/E158/D161 | GGGGS GGGGS | Venus |

**Supplementary Table 2.** Details of the sequences, manually assigned phenotypes, and phenotypes penetrances of the 1pok and 1m3u mutants. Additionally, it shows the summed value of the mutated residues of each mutant for the 20 physicochemical features analyzed in this work. (The table is available as a separated excel file)

**Supplementary Table 3.** Details of additional eight homomers targeted for alanine mutations: PDB code, number of subunits, symmetry, ORF, mutated positions, linker, and fluorescent protein fused.

| PDB code | Gene Name | Ref. | # subunits | MW (kDa, per subunit) | Symmetry | pI (WT) | pI Ala mutant | # + charges | # - charges | ORF | Mutated residues | Linker | Fluorophore |
|---|---|---|---|---|---|---|---|---|---|---|---|---|---|
| 2AN9 | *gmk* | (4) | 6 | 23.6 | D3 | 6.06 | 6.44 | 25 | 29 | 2an9-Venus(YFP) | D60/E61/K63/E64 | GGGGS GGGGS | Venus |
| 1D7A | *purE* | (5) | 8 | 16.9 | D4 | 5.82 | 6.30 | 12 | 16 | Venus-1d7a | K11/E22/E25/D158 | GGGGS GGGGS | Venus |
| 1FRW | *mobA* | (6) | 8 | 21.6 | D4 | 5.88 | 6.23 | 19 | 24 | 1frw-Venus | D170/D173/K175/D176 | GGGGS GGGGS | Venus |
| 2CG4 | *asnC* | (7) | 8 | 17 | D4 | 5.91 | 5.90 | 16 | 18 | Venus-2cg4 | K126/D131 | GGGGS GGGGS | Venus |
| 1YAC | *ycaC* | (8) | 10 | 22.9 | D5 | 5.35 | 5.32 | 20 | 25 | 1yac-Venus | D92/E94/K98/K101 | GGGGS GGGGS | Venus |
| 1L6W | *fsaA* | (9) | 10 | 23 | D5 | 5.90 | 5.55 | 17 | 19 | 1l6w-Venus | K97/K100/E102 | GGGGS GGGGS | Venus |
| 2IV1 | *cynS* | Not Published | 10 | 17 | D5 | 4.85 | 4.71 | 17 | 22 | 2iv1-Venus | K24/K25/D26 | GGGGS GGGGS | Venus |
| 2VYC | *adiA* | (10) | 10 | 84.4 | D5 | 5.12 | 5.15 | 62 | 96 | 2vyc-Venus | K491/D494/D497 | GGGGS GGGGS | Venus |

**Supplementary Table 4.** The 20 numerical scales representing physicochemical and biochemical properties of amino acids, along with their identifier in the AA index database (http://www.genome.jp/aaindex/) if applicable. (The table is available as a separated excel file)

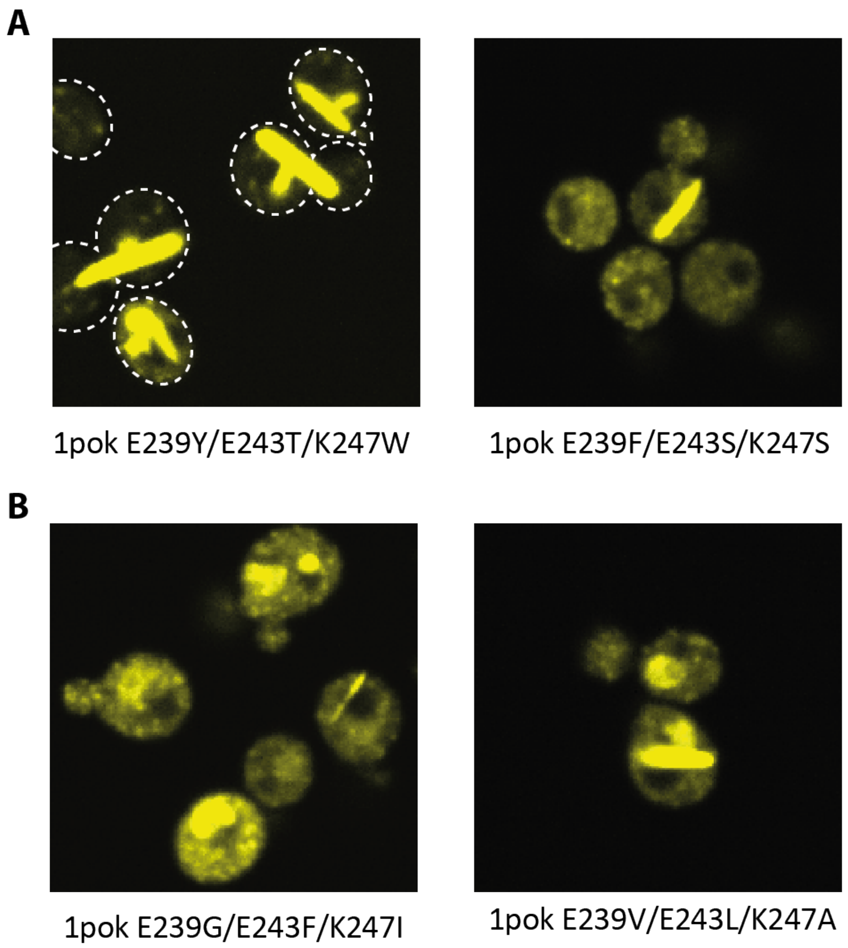


**Figure S1. Illustrative examples of phenotype heterogeneity. A** The penetrance of phenotypes can be variable, as illustrated with a mutant exhibiting fibers in most cells (left) and another mutant exhibiting fibers in a smaller fraction of cells (right). **B** Different phenotypes can also co-exist in the same cells. Both mutants show a mixture of nuclear localization and puncta phenotypes, with one cell also containing a fiber.

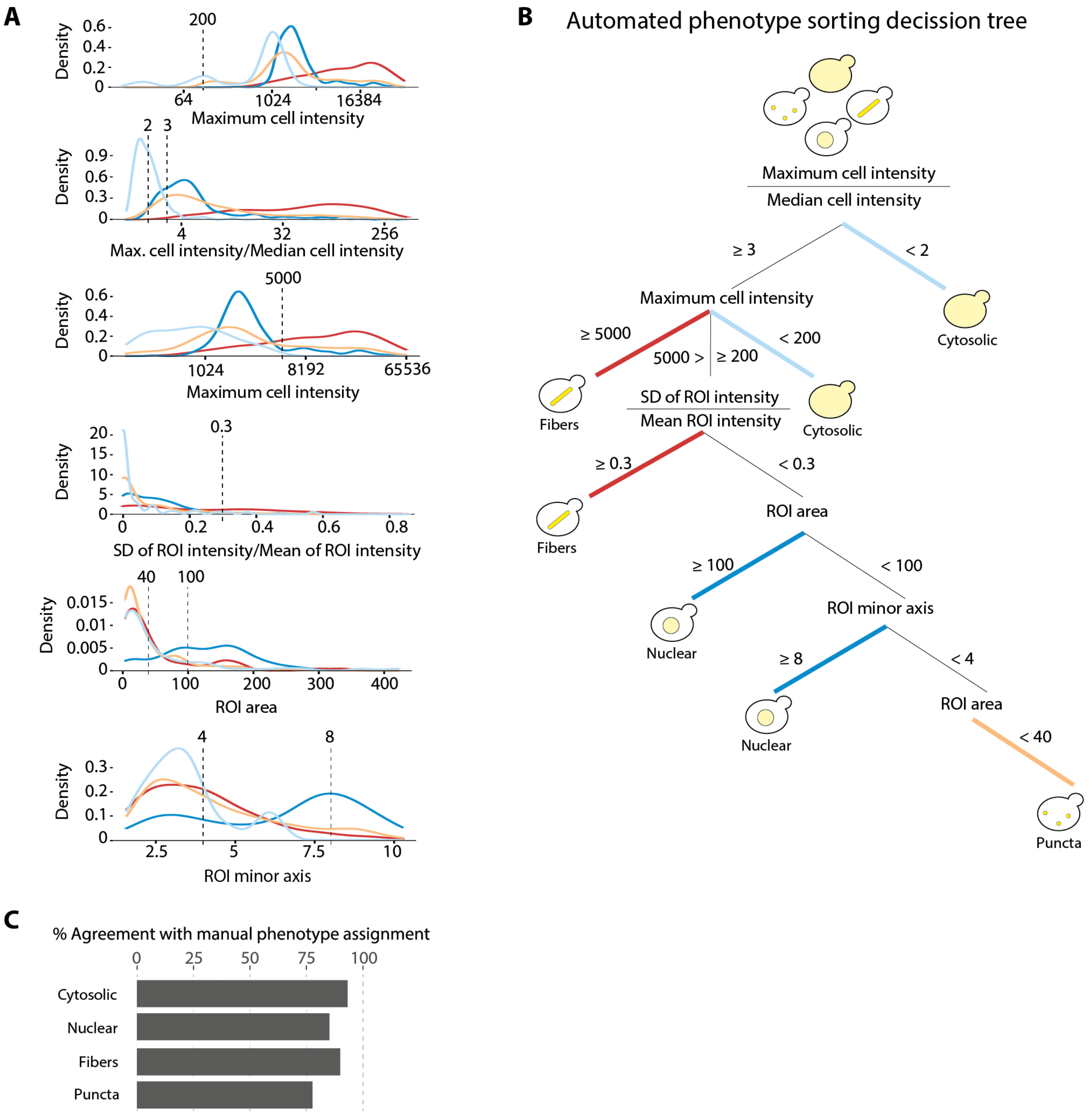


**Figure S2**. **Automated phenotype sorting. Calibration, decision tree, and validation**. **A.** Density distribution of several image properties used in the automated assignment of cells' phenotypes (Fiber, Cytosol, Nuclear, and Puncta). Dashed lines show thresholds used in the phenotype sorting decision tree. **B.** Decision tree used to classify cells and regions of interest (ROI) inside of cells. A cell is initially classified based on the maximal fluorescence intensity detected within its boundaries. Subsequently, if bright regions (maximum/median fluorescence intensity ratio > 2.5) are detected, each region is analyzed and assigned a "fiber," "nuclear," or "puncta" phenotype based on properties described in the tree. Ultimately, a cell is assigned the union of all phenotypes matching the ROIs within it. **C.** The parameters were optimized against a dataset of manually annotated cells (Methods), and we show the percent agreement between automated and manual annotations for each phenotype.

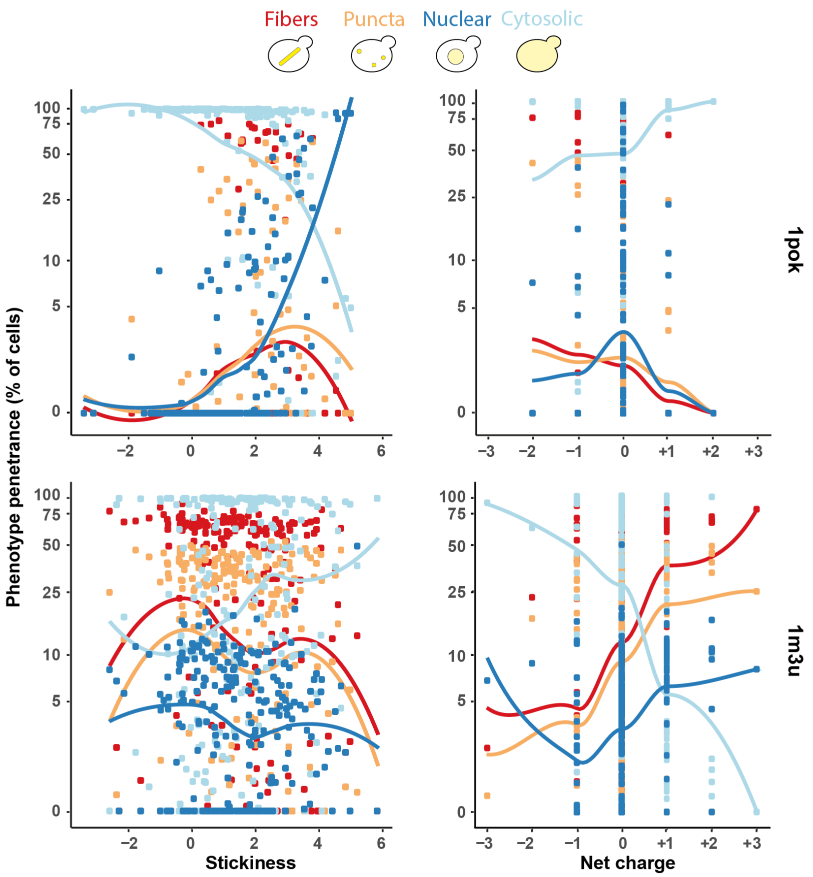


**Figure S3**. **Dependence of phenotype penetrance with the summed stickiness and net charge of the mutated triplets (all data shown)**. In the graphs, each mutant is represented by four points accounting for the fiber, puncta, nuclear and cytosolic penetrance observed in that mutant.

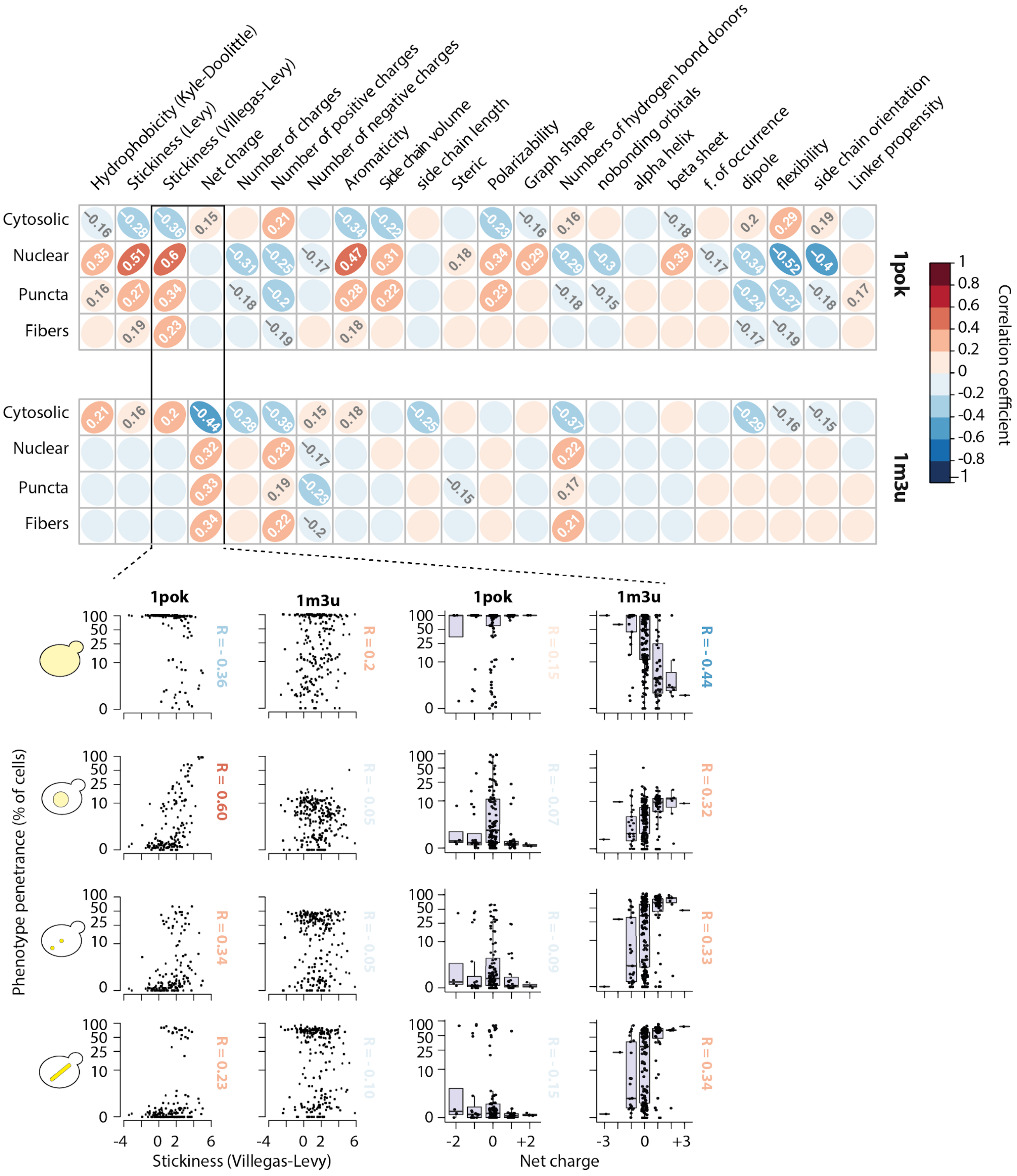


**Figure S4. Pearson correlations between phenotype penetrance and 22 physicochemical features of the three mutations observed in 1pok and 1m3u**. Only correlation coefficients larger than |0.15| are displayed. Underneath we show scatterplots of the four main phenotypes' penetrance as a function of stickiness and net charge for 1pok and 1m3u mutants.

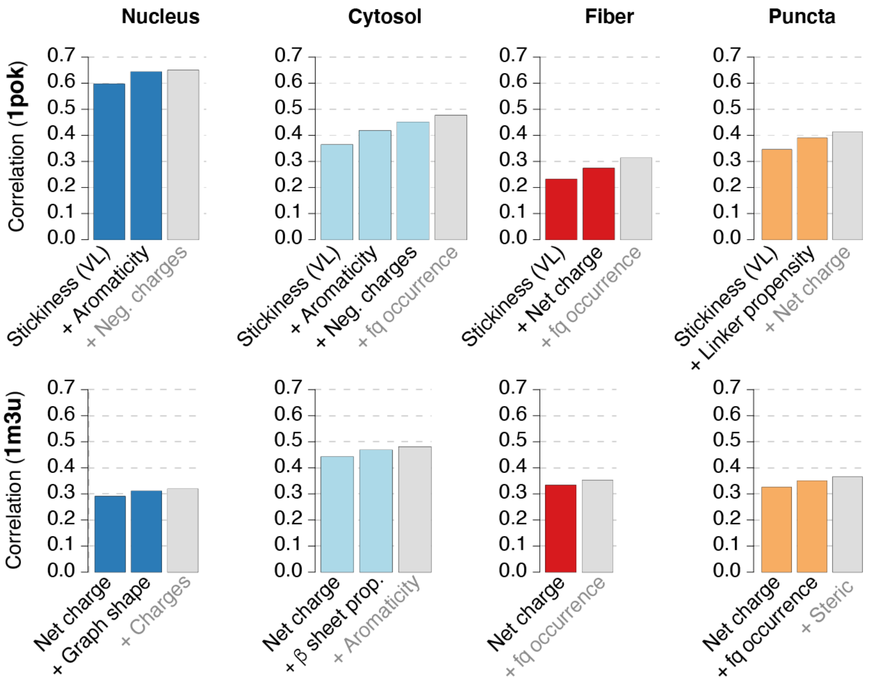


**Figure S5. Identifying physicochemical features most strongly associated with each phenotype.** For each phenotype, we selected features iteratively. The first feature was the one maximizing the correlation between phenotype penetrance and one of the 22 properties calculated of the mutated amino acids. The second feature was maximizing the partial correlation given the first feature, the third feature was maximizing the partial correlation given the first and second, etc. Up to four features were added in this way, or the procedure was stopped when a partial correlation was not significant (ANOVA, $p>0.05$). The features identified as significant are shown in color and the first non-significant feature is shown as a grey bar. In all cases, a single feature (stickiness in the case of 1pok, and net charge in the case of 1m3u) captures most of the variance that can be explained by the physicochemical properties of the mutations.

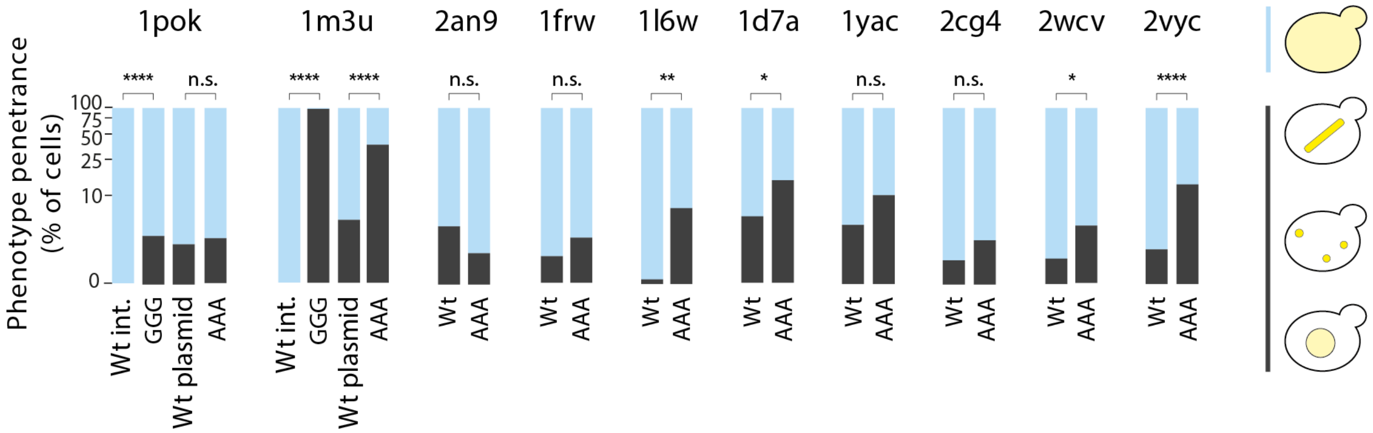


**Figure S6**. **Penetrance of non-cytosolic phenotypes for wild-type proteins and alanine or glycine mutants**. The fraction of cells with a non-cytosolic phenotype for the wild-type complex as well as alanine mutants for all ten complexes studied. Additionally, triple glycine mutants are shown for 1pok and 1m3u and compared to their genome-integrated wild-type counterparts. Statistical significance among wild-type/mutant pairs was calculated by the two-proportions Z-test.

### REFERENCES


1. D. Jozic, J. T. Kaiser, R. Huber, W. Bode, K. Maskos, X-ray structure of isoaspartyl dipeptidase from E.coli: a dinuclear zinc peptidase evolved from amidohydrolases. *J. Mol. Biol.* **332**, 243–256 (2003).
2. T. Nagai, *et al.*, A variant of yellow fluorescent protein with fast and efficient maturation for cell-biological applications. *Nat. Biotechnol.* **20**, 87–90 (2002).
3. F. von Delft, *et al.*, Structure of E. coli ketopantoate hydroxymethyl transferase complexed with ketopantoate and Mg2+, solved by locating 160 selenomethionine sites. *Structure* **11**, 985–996 (2003).
4. G. Hible, *et al.*, Calorimetric and crystallographic analysis of the oligomeric structure of Escherichia coli GMP kinase. *J. Mol. Biol.* **352**, 1044–1059 (2005).
5. I. I. Mathews, T. J. Kappock, J. Stubbe, S. E. Ealick, Crystal structure of Escherichia coli PurE, an unusual mutase in the purine biosynthetic pathway. *Structure* **7**, 1395–1406 (1999).
6. M. W. Lake, C. A. Temple, K. V. Rajagopalan, The Crystal Structure of the Escherichia coliMobA Protein Provides Insight into Molybdopterin Guanine Dinucleotide Biosynthesis. *Journal of Biological* (2000).
7. P. Thaw, *et al.*, Structural insight into gene transcriptional regulation and effector binding by the Lrp/AsnC family. *Nucleic Acids Res.* **34**, 1439–1449 (2006).
8. C. Colovos, D. Cascio, T. O. Yeates, The 1.8 A crystal structure of the ycaC gene product from Escherichia coli reveals an octameric hydrolase of unknown specificity. *Structure* **6**, 1329–1337 (1998).
9. S. Thorell, M. Schürmann, G. A. Sprenger, G. Schneider, Crystal Structure of Decameric Fructose-6-Phosphate Aldolase from Escherichia coli Reveals Inter-subunit Helix Swapping as a Structural Basis for Assembly Differences in the Transaldolase Family. *Journal of Molecular Biology* **319**, 161–171 (2002).
10. J. Andréll, *et al.*, Crystal structure of the acid-induced arginine decarboxylase from Escherichia coli: reversible decamer assembly controls enzyme activity. *Biochemistry* **48**, 3915–3927 (2009).
